# Re-examining the lower speed boundary of preferred coordination ratio constancy: estimator dependence and competing breakpoint regions

**DOI:** 10.64898/2026.04.26.720900

**Authors:** Taichi Kurayama

**Author notes:** Note on this version. This version supersedes the earlier versions of this preprint. Those versions analysed 84 individual-level coordinates recovered from panel A of the source figure. The present version analyses 95 coordinates recovered independently from panel B, where the two plotted axes are primary measured quantities on comparable linear scales and marker centres can be separated; speed and the ratio are derived per marker from the plotted cadence and step length. The change is consequential: under the earlier coordinates the breakpoint profile likelihood appeared singly peaked and a point estimate with an interval was reported, whereas under the re-digitised coordinates the profile is multimodal, with two separated regions of support near 50 and near 62 m·min⁻¹, so no single revised boundary is identified. The value reported in the source article is not refuted. The computations in the earlier versions were verified independently and were arithmetically correct; what changed is the data they were computed from. The superseded coordinate set is retained in the deposited archive alongside the current one so that the two can be compared directly. The quantity called the preferred step ratio (PSR) in earlier versions is called the preferred coordination ratio (PCR) here; the calculation is unchanged.

## Abstract

**Background:** The preferred coordination ratio (PCR), defined as step length divided by cadence, is approximately constant across a broad range of walking speeds in healthy adults, except at the slowest speeds. A previous study reported its lower boundary at approximately 62 m·min⁻¹ from K-means clustering of speed and PCR. However, K-means partitions observations rather than formally estimating a changepoint, so it is unclear how well this value identifies where PCR changes.

**Aim:** To re-examine the location and identifiability of this boundary using published data.

**Method:** Coordinates were independently recovered from the published figure, yielding speed and PCR for 95 markers. The reported K-means-derived boundary was reproduced and compared with changepoint and segmented-model estimates. Sensitivity analyses examined digitisation and model specification.

**Results:** For the reference robust heteroscedastic segmented model, the profile likelihood was multimodal, with a global maximum at 50 m·min⁻¹ and a competing mode at 62 m·min⁻¹; the twice-log-likelihood difference was 0.82. The support set was disconnected: 43–52 and 56–66 m·min⁻¹. Across eight segmented-model specifications the maximising breakpoint ranged from 45 to 62 m·min⁻¹, while a Gaussian mean-and-variance changepoint estimator yielded 38 m·min⁻¹. The reference segmented estimate remained at 50 m·min⁻¹ in all coordinate-recovery analyses. The published K-means-derived boundary was reproduced at 62.4 m·min⁻¹.

**Interpretation:** The reanalysis found competing breakpoint regions rather than a single dominant boundary. Alternative formulations indicated that the boundary may lie below approximately 62 m·min⁻¹, warranting direct testing with new data.

**Highlights:**

- Formal analyses found competing regions rather than one dominant breakpoint
- The reference profile peaked at 50 m/min; a competing mode occurred at 62 m/min
- Several analyses indicated that the transition may occur below 62 m/min

## 1. Introduction

The ratio of step length to cadence has long been reported to be approximately constant across a broad range of walking speeds in healthy adults ^1,2^. In recent studies, this ratio has been called the walk ratio (WR) ^2,3,4^. In this study, the same quantity is called the preferred coordination ratio (PCR) to emphasise the freely selected coordination of step length and cadence. In community-dwelling older adults, the ratio was similar during preferred-speed, fast, and uneven-surface walking but changed during a concurrent cognitive task ^3^. Departures from the freely selected ratio have been reported to increase the mechanical power required for walking ^5^, suggesting that such departures carry a biomechanical cost. Sekiya et al. ^2^ reported that the ratio remained relatively constant above approximately 60 m·min⁻¹ but was higher and more variable at lower speeds.

Murakami and Otaka ^4^ applied two-cluster K-means clustering to speed and PCR and reported a non-overlapping separation between slow- and normal-walking observations at approximately 62 m·min⁻¹ (≈1.03 m·s⁻¹). Their study is important in that it treated the location of the change as the object of an explicit clustering analysis.

However, the exact numerical location of that boundary remains open to examination. K-means partitions observations by minimising within-cluster squared distances; it is not a changepoint method that directly estimates where the distribution of PCR changes. The boundary defined from that separation also depends on the observations adjacent to it on both sides. In contrast, changepoint and segmented-regression methods directly estimate where a specified feature of the response distribution changes ^6,7^.

This study re-examined the statistical location and identifiability of the lower boundary of PCR constancy using the published data of Murakami and Otaka ^4^. The source figure was independently re-digitised, and the recovered coordinates were analysed using changepoint and segmented-regression methods. To assess the robustness of the boundary estimate, sensitivity to estimator choice, model specification, and coordinate recovery was examined. The published K-means-derived boundary was also reproduced as a separate comparison.

## 2. Methods

The segmented model was used as the reference formal analysis. Sensitivity analyses examined estimator choice, model specification, and coordinate recovery. The published K-means result was reproduced as a separate comparison. The separate-breakpoint analysis was exploratory.

### 2.1 Source study and published information

This reanalysis used only information available in the published report of Murakami and Otaka ^4^. The analysis used the within-cluster Pearson correlations between speed and PCR; the condition-specific cluster counts reported in their Table 1; the maximum speed of the slow-walk cluster and the minimum speed of the normal-walk cluster reported in their Results; and their Figure 1.

**Figure 1.**
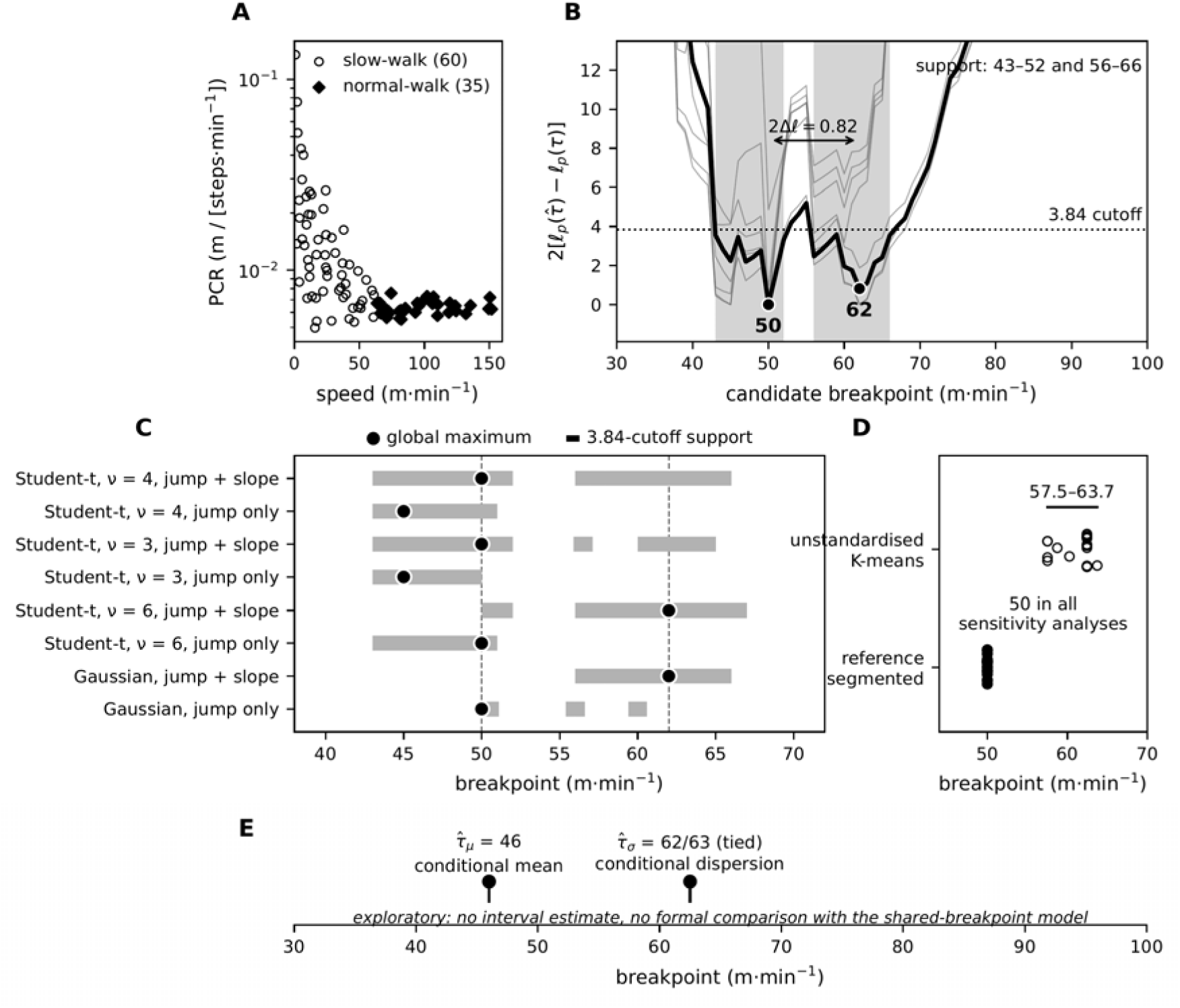
Reanalysis of the lower speed boundary of preferred coordination ratio (PCR) constancy. **(A)** The 95 markers recovered from panel B of the source figure: 60 in the slow-walk cluster (open circles) and 35 in the normal-walk cluster (filled diamonds). Speed and PCR were calculated for each marker from the plotted cadence and step length. The vertical axis is logarithmic. **(B)** Profile likelihood of the reference segmented model (Student-t, ν = 4; dispersion jump and slope), heavy line, with the seven other specifications in light grey. The dotted line marks the 3.84 cutoff; shading marks the disconnected support set. The global maximum is at 50 m·min⁻¹ and the competing higher-speed mode at 62 m·min⁻¹, with a twice-log-likelihood difference of 0.82. **(C)** Model-specification sensitivity: for each specification, the global maximum (filled circle) and the support set (bars). Dashed guides at 50 and 62 m·min⁻¹. **(D)** Sensitivity to coordinate recovery. Across the coordinate-recovery analyses, the unstandardised K-means boundary ranged from 57.5 to 63.7 m·min⁻¹, whereas the breakpoint from the reference segmented model remained at 50 m·min⁻¹. **(E)** Exploratory model with separate breakpoints for the mean and the dispersion. The mean breakpoint was 46 m·min⁻¹; the dispersion breakpoint showed equal maxima at 62 and 63 m·min⁻¹, which could not be distinguished. No interval was estimated, and no formal comparison with the shared-breakpoint model was made. All support sets were calculated under the working-independence assumption; no nominal coverage is claimed.

**Table 1.** Direct estimates from the primary reconstructed coordinates (n = 95). Support sets are profile-likelihood sets defined by the conventional 3.84 cutoff under a working-independence representation; no nominal coverage is claimed.

| Estimator | Variables analysed | Estimate, $\text{m} \cdot \text{min}^{-1}$ | $\text{m} \cdot \text{s}^{-1}$ | Support set, $\text{m} \cdot \text{min}^{-1}$ |
| --- | --- | --- | --- | --- |
| Unstandardised K-means<br>(reconstruction of the published analysis) | speed, PCR | 62.4 | 1.04 | — |
| Standardised K-means | speed, PCR (z) | 57.5 | 0.96 | — |
| Gaussian mean-and-variance changepoint | raw PCR | 38 | 0.63 | — |
| Reference segmented specification, global maximum | log PCR | 50 | 0.83 | 43–52, 56–66 |
| Reference segmented specification, competing mode | log PCR | 62 | 1.03 | (within the same support set) |

*Note. Under the reference specification, $2[\ell_{\square}(50) - \ell_{\square}(62)] = 0.82$ .*

No new participant data were collected, and the original individual-level data were not available to this study. Because the analysis used only published aggregate information and coordinates recovered from the published figure, institutional ethics review was not required.

### 2.2 Independent re-digitisation

The source figure was extracted from the publisher PDF at native resolution. Marker coordinates were recovered from panel B (adjusted step length versus adjusted cadence) because the markers were more clearly separated there. Pixel coordinates were converted to step length and cadence by automatically detecting axis ticks and fitting linear calibration functions between pixel position and tick value. Speed and PCR were then calculated for each marker as follows: speed = cadence × step length and PCR = step length/cadence. Figure extraction, calibration, and coordinate recovery were implemented in Python; software versions and full reproducibility details are provided in section 2.10.

### 2.3 Marker detection and audit

Markers were detected automatically by connected-component image analysis and assigned to the two clusters indicated by the source-study symbols. Detection parameters were fixed before the statistical analysis and were not chosen to match the published statistics. The recovered coordinates were checked visually against the source figure. No manual corrections were made. Panel B represented 105 observations (21 participants × 5 conditions) and yielded 103 detectable components. Of these, 95 markers were fully resolved and formed the primary reconstructed dataset. Six partially resolved merged blobs, all from the normal cluster, could not be assigned uniquely to individual markers and were excluded without coordinate imputation. Two unresolved overlaps were excluded from the primary dataset and examined using six treatments, including exclusion as the reference treatment (section 2.8). Two observations yielded no separable component. Detection and audit details are in the deposited archive.

### 2.4 Reproduction of the published K-means-derived boundary

Two-cluster K-means was applied to the recovered speed and PCR coordinates, to assess whether the published boundary could be reproduced. The source study did not report standardising speed or PCR. The reported analysis was therefore interpreted as using the original scales and is referred to below as unstandardised K-means. The clustering was taken as the partition minimising the within-cluster sum of squares. An optimal two-means partition is separated by a hyperplane. The optimum was therefore obtained by projecting the observations onto the direction defined by each pair of points, splitting each resulting sorted order, and enumerating the resulting partitions ^8^, rather than by restarted local search. The boundary was then taken as the midpoint between the fastest observation assigned to the slow-walk cluster (61.1 m·min⁻¹) and the slowest assigned to the normal-walk cluster (63.6 m·min⁻¹). The source study reported these two speeds and stated that the clusters divided at approximately 62 m·min⁻¹ without overlap. Its Figure 1A marked the boundary at 62.4 m·min⁻¹, but the method used to obtain this value was not stated. The midpoint is the reading adopted here, and it reproduces the value printed in the figure. The analysis was then repeated after z-standardising both variables, to assess sensitivity to feature scaling.

### 2.5 Gaussian mean-and-variance changepoint estimator

As an estimator-sensitivity analysis, a Gaussian changepoint model examined an abrupt change in the mean and variance of raw PCR.

Candidate breakpoints from 5 to 100 m·min⁻¹ were evaluated at 1 m·min⁻¹ intervals. At each candidate, the observations were divided into a lower- and a higher-speed segment, each required to contain at least five observations, and a Gaussian model allowing separate means and separate variances in the two segments was compared with a single-segment Gaussian model by a generalised likelihood-ratio statistic ^9^. The candidate maximising that statistic was retained as the estimate.

### 2.6 Robust heteroscedastic segmented model

The reference formal analysis modelled the low-speed transition in log-PCR. Above a candidate breakpoint, mean log-PCR was held constant. Below it, mean log-PCR was allowed to rise as speed fell. Dispersion was allowed to change at the breakpoint and to increase further at lower speeds. A Student-t distribution with ν = 4 was used to limit the influence of individual extreme observations. Together with the full dispersion model, this formed the reference specification ^10^; other specifications are compared in section 2.7.

Let y_i_ = log(PCR_i_), v_i_ = speed_i_, and h_i_(τ) = max(τ − v_i_, 0). For each candidate breakpoint τ on the same 5–100 m·min⁻¹ grid at 1 m·min⁻¹ intervals,

- y_i_ ∼ t_ν_(μ_i_, σ_i_);
- μ_i_ = α + β h_i_(τ), β ≥ 0;
- log σ_i_² = γ₀ + γ₁·1(v_i_ < τ) + γ₂ h_i_(τ), γ₁, γ₂ ≥ 0.

No minimum number of observations was required on either side of a candidate; every maximising breakpoint and the support set lay well inside the grid. At each τ the remaining parameters were estimated by constrained maximum likelihood, and the τ maximising the profile log-likelihood was taken as the estimate ^7^.

The entire profile likelihood was evaluated to examine support for other candidates. As a descriptive criterion, the conventional cutoff of 3.84 (the 95th percentile of a χ² distribution with one degree of freedom) was used to define a profile-likelihood support set, {τ: 2[ℓₚ(τ^) − ℓₚ(τ)] ≤ 3.84}. The set was reported in its observed form, including when it was disconnected. Breakpoint inference is non-regular ^11,12^. This cutoff has not been calibrated for the present design, so the set is not interpreted as a 95% confidence interval and no nominal coverage is claimed for it.

The estimator of section 2.5 uses raw PCR and a two-segment Gaussian model, whereas this section uses log-PCR and a segmented model; the two are different statistical formulations, and their numerical estimates should not be interpreted as the same parameter on different scales. Because the published figure did not permit observations across the five speed conditions to be linked within participants, within-participant dependence could not be modelled. The likelihood analyses therefore used a working-independence specification ^13^.

### 2.7 Model-specification sensitivity

To assess whether the breakpoint estimate depended on the statistical specification, the same primary reconstructed coordinates were analysed under eight segmented-model specifications. The eight specifications combined four error distributions (Student-t with ν = 3, 4 or 6, and Gaussian) with two dispersion structures: the full model, allowing both a dispersion jump and a below-breakpoint dispersion gradient, and a simplified model with the gradient removed (γ₂ = 0). The complete breakpoint profile was computed for each, and a support set was constructed under the same 3.84 criterion as in section 2.6.

### 2.8 Digitisation sensitivity analyses

Sensitivity to coordinate recovery was assessed in two series. The series were evaluated separately and were not crossed. The first examined 13 perturbed marker-detection settings; one was excluded because its peak threshold recovered no normal-walk markers at all, leaving 12. The second evaluated six treatments of the two unresolved components described in section 2.3: the primary exclusion and five alternatives. In both series, the unstandardised and standardised K-means boundaries, the mean-and-variance changepoint estimate, and the segmented breakpoint were recomputed. The segmented model was kept at its reference specification throughout.

### 2.9 Exploratory two-breakpoint model

To examine whether the mean and dispersion components of the segmented model might change at different speeds, an exploratory variant allowed separate breakpoints for the conditional mean (τ_μ_) and the conditional dispersion (τ_σ_):

- μ_i_ = α + β h_i_(τ_μ_), β ≥ 0;
- log σ_i_² = γ₀ + γ₁·1(v_i_ < τ_σ_) + γ₂ h_i_(τ_σ_), γ₁, γ₂ ≥ 0,

with Student-t errors at ν = 4, as in the reference specification. All pairs of breakpoints on the same 5–100 m·min⁻¹ grid at 1 m·min⁻¹ intervals were fitted. Each segment was required to contain at least three observations. No interval was estimated, and the model was not formally compared with the shared-breakpoint model.

### 2.10 Reproducibility and software

Figure calibration, marker recovery, and all primary statistical analyses were implemented in Python. A separate MATLAB implementation was used only for cross-language validation. It reproduced the corresponding direct estimates and breakpoint-profile structure. A single Python entry point regenerates all reported results from the deposited coordinates and writes machine-readable outputs. The archive includes the Python and library versions, a version-fixed requirements file, and the figure-extraction and axis-calibration diagnostics.

## 3. Results

### 3.1 Recovered data and source consistency

The primary reconstructed dataset comprised 95 markers (Figure 1A). Several quantities agreed closely with values reported in the source article: the maximum speed in the slow-walk cluster was 61.21 m·min⁻¹ (reported, 61.1), the minimum speed in the normal-walk cluster was 63.67 m·min⁻¹ (reported, 63.6), and the speed–PCR correlation within the slow-walk cluster was −0.4655 (reported, −0.47).

The corresponding correlation within the normal-walk cluster was +0.1602 rather than the reported +0.03. In addition, at least 60 markers were assigned to the slow-walk cluster in the source figure, whereas the condition-specific counts in the source article’s Table 1 sum to 44 observations in that cluster. These discrepancies could not be resolved from the publicly available information.

### 3.2 Reproduction and sensitivity of the published K-means-derived boundary

Using this reconstruction, the K-means-derived boundary was 62.4 m·min⁻¹ (1.04 m·s⁻¹) on the primary reconstructed coordinates, matching the published value (Table 1). The standardised variant returned 57.5 m·min⁻¹.

Across the 12 retained perturbed detection settings and the six treatments of the unresolved components, the unstandardised K-means-derived boundary ranged from 57.5 to 63.7 m·min⁻¹ (0.96 to 1.06 m·s⁻¹), and the standardised variant from 49.0 to 57.5 m·min⁻¹ (Table 2, Figure 1D).

**Table 2.** Sensitivity of the estimates to coordinate recovery, with the reference model specification fixed. Analyses included 12 retained perturbed detection settings and six treatments of the two unresolved image components.

| Estimator | Baseline | Across 12 detection settings | Across 6 treatments |
| --- | --- | --- | --- |
| Reference segmented specification, maximising breakpoint | 50 | 50 in all | 50 in all |
| Gaussian mean-and-variance changepoint | 38 | 38 in all | 38 or 43 |
| Unstandardised K-means-derived boundary | 62.4 | 57.5 to 63.7 | 57.5 to 62.4 |
| Standardised K-means | 57.5 | 49.0 to 57.5 | 49.0 to 57.5 |

*Note. A thirteenth detection setting was excluded for structural reasons. At a peak threshold of 0.40, no filled diamonds were* *recovered because a diamond with the observed geometry has a maximum distance transform of approximately 0.40 times its width.* *Thus, this threshold removed the marker class by construction rather than because of the data.*

### 3.3 Profile-likelihood structure of the segmented model

Under the reference specification (Student-t, ν = 4, full dispersion model), the profile likelihood over the breakpoint was multimodal (Figure 1B). Within the profile-likelihood support set, local maxima occurred at 45, 47, 50, 56 and 62 m·min⁻¹, forming two separated regions of support. The global maximum occurred at τ^ = 50 m·min⁻¹ (0.83 m·s⁻¹). A competing higher-speed maximum occurred at 62 m·min⁻¹, at which

2[ℓₚ(50) − ℓₚ(62)] = 0.82,

below the 3.84 cutoff, so both 50 and 62 m·min⁻¹ belonged to the support set.

The 3.84-cutoff profile-likelihood support set was disconnected, comprising 43–52 and 56–66 m·min⁻¹; the region from 53 to 55 m·min⁻¹ did not satisfy the cutoff.

### 3.4 Model-specification sensitivity

Refitting the same coordinates under eight specifications yielded maximising breakpoints ranging from 45 to 62 m·min⁻¹ (Table 3, Figure 1C). The full-dispersion models yielded 50 m·min⁻¹ with Student-t errors (ν = 3 or 4) and 62 m·min⁻¹ with Student-t errors (ν = 6) or Gaussian errors. Removing the below-breakpoint dispersion gradient yielded 45 or 50 m·min⁻¹.

**Table 3.** Model-specification sensitivity of the segmented model, fitted to the same primary reconstructed coordinates. The maximising breakpoint ranged from 45 to 62 m·min⁻¹. All eight specifications had local maxima in the lower region (43–53) and the higher region (56–67); the region containing the global maximum depended on the specification.

| Error distribution | Dispersion structure | Maximising breakpoint, $\text{m} \cdot \text{min}^{-1}$ | 3.84-cutoff support set, $\text{m} \cdot \text{min}^{-1}$ |
| --- | --- | --- | --- |
| Student-t, $\nu = 4$ | jump and slope | 50 | 43–52, 56–66 |
| Student-t, $\nu = 3$ | jump and slope | 50 | 43–52, 56–57, 60–65 |
| Student-t, $\nu = 6$ | jump and slope | 62 | 50–52, 56–67 |
| Gaussian | jump and slope | 62 | 56–66 |
| Student-t, $\nu = 4$ | jump only ($\gamma = 0$) | 45 | 43–51 |
| Student-t, $\nu = 3$ | jump only ($\gamma = 0$) | 45 | 43–50 |
| Student-t, $\nu = 6$ | jump only ($\gamma = 0$) | 50 | 43–51 |
| Gaussian | jump only ($\gamma = 0$) | 50 | 50–51, 56, 60 |

*Note. The first row is the reference specification. The support set was disconnected in four of the eight specifications.*

The support set was disconnected in four of the eight specifications.

Local maxima were present in both the lower-speed region (43–53 m·min⁻¹) and the higher-speed region (56–67 m·min⁻¹) under all eight specifications. The region containing the global maximum depended on the specification.

### 3.5 Gaussian mean-and-variance changepoint estimate

The Gaussian mean-and-variance changepoint estimator on raw PCR returned 38 m·min⁻¹ (0.63 m·s⁻¹) (Table 1).

### 3.6 Digitisation sensitivity

With the reference specification held fixed, the global maximum of the segmented model was 50 m·min⁻¹ in all 12 retained perturbed detection settings and all six treatments. The mean-and-variance changepoint estimate was 38 m·min⁻¹ under all 12 detection settings; under the six treatments it was 38 m·min⁻¹ in three and 43 m·min⁻¹ in three (Table 2, Figure 1D).

### 3.7 Exploratory separate mean and dispersion breakpoints

The mean transition was estimated at τ^_μ_ = 46 m·min⁻¹, whereas the dispersion transition had tied maxima at 62 and 63 m·min⁻¹ (Figure 1E). No observations lay between 61.21 and 63.67 m·min⁻¹, so the data could not distinguish the two maxima. This exploratory analysis did not test whether the mean and dispersion breakpoints differed.

## 4. Discussion

### 4.1 Main findings

When the changepoint was modelled directly, the profile likelihood was multimodal. Its global maximum occurred at 50 m·min⁻¹, with a competing mode at 62 m·min⁻¹ and a disconnected support set. The maximising breakpoint varied from 45 to 62 m·min⁻¹ across specifications, and the Gaussian mean-and-variance changepoint estimator yielded 38 m·min⁻¹. The estimated boundary therefore depended on the statistical formulation: six of the eight segmented specifications and the changepoint estimator placed it below the published value.

Using the unstandardised K-means analysis reconstructed from the source study ^4^, the published boundary was reproduced at 62.4 m·min⁻¹. The evidence that PCR changes at slow walking speeds remains intact, and the present reanalysis does not challenge this broader finding. The previously reported value of approximately 62 m·min⁻¹ also remains a supported candidate: it lay within the support set under the reference and the other full-dispersion specifications, and a local maximum occurred near it under all eight.

The results do not justify replacing 62 m·min⁻¹ with 50 m·min⁻¹, 38 m·min⁻¹, or another single revised value. The analyses did not determine a single alternative boundary.

### 4.2 Why the K-means-derived boundary and formal breakpoint estimates differ

K-means partitions the observations, whereas the formal breakpoint methods estimate where the distribution changes. The reported value lies within the gap between the two clusters’ adjacent extreme observations. The width and location of this gap depend on the observations immediately adjacent to the separation. When the midpoint is used, a change in either observation moves the boundary. In the unstandardised reconstruction, differences on the numerically larger speed scale contributed disproportionately to the K-means clustering ^14^. Consistent with this, the standardised variant returned 57.5 m·min⁻¹ (49.0–57.5 across coordinate-recovery conditions).

In the reanalysis the K-means boundary moved between 57.5 and 63.7 m·min⁻¹ across coordinate-recovery conditions. The reference segmented breakpoint, by contrast, was 50 m·min⁻¹ under every such condition. Within the coordinate-recovery conditions examined, the estimate of 50 m·min⁻¹ did not depend on the digitisation. However, changing the statistical model on the same coordinates moved the estimate between 45 and 62 m·min⁻¹, so it cannot be regarded as a uniquely determined boundary.

### 4.3 Transitions below the published boundary and their structure

Two different formulations yielded point estimates below approximately 62 m·min⁻¹: the reference segmented model was maximised at 50 m·min⁻¹, and the Gaussian mean-and-variance changepoint estimator at 38 m·min⁻¹ (43 m·min⁻¹ under three of the six treatments of the unresolved components). The exploratory model allowing separate mean and dispersion breakpoints likewise placed the mean transition at 46 m·min⁻¹. These estimates come from different statistical formulations and should not be treated as repeated estimates of the same parameter. However, alternative formulations placed the transition below the published value, providing a reason to examine lower speeds directly.

Replacing the published boundary with another single value may also be insufficient. In the exploratory two-breakpoint analysis, the breakpoint for the conditional mean was 46 m·min⁻¹, whereas that for the conditional dispersion was tied at 62 and 63 m·min⁻¹. If mean PCR and its variability change at different speeds, the speed dependence of PCR need not be a single boundary dividing a higher-from a lower-speed regime, and may involve more than one transition. Sekiya et al. ^2^ reported both the rise and the increased variability below about 60 m·min⁻¹, consistent with this reading, but did not separate the two locations. Because the exploratory model provided no uncertainty interval and was not formally compared with the shared-breakpoint model, this interpretation remains a hypothesis.

Several analyses indicated that the transition relevant to PCR constancy may occur below approximately 62 m·min⁻¹. Studies with denser repeated measurements are needed to determine whether the mean and variability of PCR change at the same or different speeds.

### 4.4 Source-data constraints

Not all reported values could be reconciled, internally or with the recovered coordinates. At least 60 markers were assigned to the slow-walk cluster in the re-digitised figure, whereas the condition-specific counts in the source article’s Table 1 sum to 44 observations in that cluster. Because the original individual-level data are unavailable, this discrepancy cannot be resolved. The correlation mismatch (+0.16 recovered versus +0.03 reported) may have the same source: the six excluded partially resolved components all belonged to the normal-walk cluster, and including them moves the recovered correlation to +0.10. The formal breakpoint estimators do not use the published cluster assignments. The count and correlation mismatches are therefore directly relevant to reproducing the published clustering. However, unrecovered markers may also have affected the breakpoint estimates. The main interpretation therefore rests on results obtained directly from the recovered coordinates.

### 4.5 Limitations and implications

This study has several limitations. First, the analyses used coordinates recovered from a published figure rather than the original individual-level data. Uncertainty in the recovery was assessed by sensitivity analysis. Ten of the 105 observations did not enter the primary dataset — six partially resolved components, all from the normal cluster; two unresolved overlaps, assessed separately in the sensitivity analyses; and two observations yielding no separable component — so selection effects cannot be ruled out.

Second, the within-subject structure of the original experiment could not be modelled because participant identities could not be recovered from the figure. The profile-likelihood support sets are conditional on this working-independence representation and should not be interpreted as population-level confidence intervals.

Third, the original study included only five prescribed speed conditions, leaving the competing breakpoint regions sparsely sampled. Denser observations around these regions are required to determine the transition more precisely.

Given these limitations, direct studies with participant-level repeated measurements are needed to determine whether the transition relevant to PCR constancy occurs at speeds below those previously reported. Such studies would clarify the speed range over which step length–cadence coordination changes.

## 5. Conclusions

The published K-means-derived boundary was reproduced at 62.4 m·min⁻¹. However, several formal analyses indicated that the transition may occur below approximately 62 m·min⁻¹. This possibility requires direct testing with individual-level data and denser speed sampling.

## Conflict of Interest

The author declares no competing financial or professional interests.

## Data Availability

The source article and its figure are freely available from the publisher. The recovered coordinate data, code and analysis outputs generated in this study are openly deposited at: Kurayama T. The lower speed boundary of preferred coordination ratio constancy: re-digitised coordinates, analysis code and results (v1.1.0) [Software]. Zenodo; 2026. https://doi.org/10.5281/zenodo.21951380. The published source figure is not redistributed. This reanalysis was not preregistered with an analysis plan in an independent registry.

## Preprint

This manuscript is posted as the corrected version of the preprint at https://doi.org/10.64898/2026.04.26.720900, which this version supersedes together with its earlier versions (see the note on this version above). It has been submitted to a peer-reviewed journal.

## Declaration of Generative AI and AI-Assisted Technologies in the Manuscript Preparation Process

During the preparation of this manuscript, the author used Anthropic Claude and OpenAI ChatGPT for analysis-code development, statistical and structural review, and editing of the manuscript, including English proofreading. The author reviewed and edited the content as necessary and takes full responsibility for the final version of the manuscript; every computational result was regenerated from the deposited code and checked against the machine-readable outputs. No AI-generated illustrations were used.

## Notes

https://doi.org/10.5281/zenodo.21951380

### Competing Interest Statement

The authors have declared no competing interest.

### Summary of Updates

This version supersedes all earlier versions, which analysed 84 coordinates recovered from panel A of the source figure. The present version analyses 95 coordinates independently recovered from panel B. With the new coordinates, the breakpoint profile likelihood is multimodal, with separate support regions near 50 and 62 m/min. No single revised boundary is therefore identified, and the originally reported value is not refuted. The superseded coordinate set is retained in the deposited archive. The quantity called the preferred step ratio (PSR) in earlier versions is now called the preferred coordination ratio (PCR); the calculation is unchanged.

